# Differential encoding of safe and risky offers

**DOI:** 10.1101/2021.01.07.425153

**Authors:** David J-N. Maisson, Seng Bum Michael Yoo, Maya Zhe Wang, Tyler V. Cash-Padgett, Jan Zimmermann, Benjamin Y. Hayden

## Abstract

Common currency theories in neuroeconomics hold that neurons in specific brain regions specifically encode subjective values of offers and not stimulus-specific information. The rationale behind these theories is that abstract value encoding lets the decision maker compare qualitatively different options. Alternatively, expectancy-based theories hold that the brain preferentially tracks the relationship between options and their outcomes, and thus does not abstract away details of offers. To adjudicate between these theories, we examined responses of neurons in six reward regions to risky and safe offers while macaques performed a gambling task. In all regions, responses to safe options are unrelated to responses evoked by equally preferred risky options. Nor does any region appear to contain a specialized subset of value-selective neurons. Finally, in all regions, responses to risky and safe options occupy distinct response subspaces, indicating that the organizational framework for encoding risky and safe offers is different. Together, these results argue against the idea that putative reward regions carry abstract value signals, and instead support the idea that these regions carry information that links specific options to their outcomes in support of a broader cognitive map.

## INTRODUCTION

The idea that the brain makes use of a common currency (sometimes called abstract) value representations has been foundational within neuroeconomics and the neuroscience of reward (Montague & Berns, 2002; FitzGerald et al., 2009; Kable & Glimcher, 2009; Padoa-Schioppa, 2011; Levy & Glimcher, 2012; Gross et. al, 2014; O’Donoghue & Rabin, 2015). Indeed, the identification of abstract value representation has been identified as the central research problem in the field (Rangel et al., 2008; Kable & Glimcher 2009; Padoa-Schioppa & Conen, 2017). A *sine qua non* of common currency coding is the existence of *pure value neurons* - neurons whose firing rate encodes the value of an offer on a single scale (Dorris & Glimcher, 2004; Padoa-Schioppa & Assad, 2006; Klein et al., 2008; Lau & Glimcher, 2008; Xie & Padoa-Schioppa, 2016). A common currency scale must by definition be amodal, meaning that responses of a value neuron to two offers must be identical if the values of the offers are the same, even if the offers differ in other ways, such as their composition or their location in space. Such abstract value representations are potentially beneficial because they allow for neutral comparison of qualitatively different goods. Consequently, they are central to several models of choice (e.g. Glimcher et al., 2005; Rustichini & Padoa-Schioppa, 2015).

The orbitofrontal cortex (OFC) has been central to debates about common currency encoding (Tremblay & Schultz, 1999; Rolls, 2000; O’Doherty et. al, 2001; Wallis, 2007; Padoa-Schioppa, 2011; Padoa-Schioppa & Schoenbaum, 2015; Wang & Hayden, 2017). While much work supports the idea that the OFC carries a common currency value representation, a complementary body of literature links OFC to a different and more general function - predicting outcomes (Schoenbaum et al., 1998; Schoenbaum et al., 2003; Kahnt et al., 2010; Schoenbaum et al., 2011; Takahashi et al., 2011; Farovik et al., 2015; Lucantonio et al., 2015). This expectancy view predicts that neuronal responses should be specific to the properties of the outcome, and not just to its value, so two qualitatively different offers with the same value will generally elicit distinct and unrelated neural responses. These ideas in turn motivated and served as the foundation for cognitive mapping theories, which hold that the brain contains specialized regions whose responses map stimuli to their expected outcomes, and, overall, implement a cognitive map of task space (Wilson et al., 2014; Schuck et al., 2016; Wickenheiser & Schoenbaum, 2016; Behrens et al., 2018; Schuck & Niv, 2019).

A good deal of evidence supports the idea that the brain contains neurons whose responses reflect value. In particular, several brain regions include neurons with statistically equivalent responses to equally-valued offers that are defined by different combinations of the same attributes (Padoa-Schioppa & Assad, 2006; Rudebeck & Murray, 2014; Strait et al., 2014; Rudebeck et al., 2017; Azab & Hayden 2020). For example, neurons in the orbitofrontal cortex (OFC, area 13) will respond the same way to two offers with, respectively, a small amount of a more preferred juice and a larger amount of a less preferred juice (Padoa-Schioppa & Assad, 2006). Likewise, neurons in the ventromedial prefrontal cortex (vmPFC, area 14) show a positive correlation between regression weights for the stakes and probability of risky offers (Strait et al., 2014). Both results reflect the same underlying process - a shedding of information (i.e. abstraction) about the details of the factors that produce the value. However, these findings are fundamentally negative ones - they identify a small range of conditions in which neurons fail to distinguish equally valued but different offers, leaving open the possibility that common currency coding is violated in the more general case of dissimilar goods.

Perhaps the most well-known and well-studied example of comparison of dissimilar goods is the choice between risky and safe options (McCoy & Platt, 2005; Platt & Huettel, 2008; So & Stupohorn, 2010; Kim et al., 2012; So & Stuphorn, 2016; Farashahi et al., 2019). Decision-makers, both in the lab and in the world, are often faced with a choice between options that offer either a guaranteed sum (e.g., $10) or the result of an unpredictable stochastic process (e.g., a 50% chance of $0 and a 50% chance of $20). Notably, risky and safe options are psychologically different in several respects - for example, the risky option may elicit a greater potential for learning or engender a different affective response (Lopes, 1987; Loewenstein et al., 2001; Barseghyan et al., 2013). Nonetheless, humans and monkeys can adroitly compare risky and safe offers to each other in an economically meaningful way (Heilbronner, 2017). Consequently, common currency models necessarily predict the existence of neurons with identical neural responses to equally valued risky and safe offers. Any observable difference between responses to these offers would make them distinguishable and thus violate the core definition of common currency value encoding (Levy & Glimcher, 2012).

Here we asked whether neural responses to risky and safe offers use a common currency value code in any of six core reward regions of the brain. Because of ongoing unresolved debates about the potential locus of abstract value encoding, we investigated six regions using the same task: the ventromedial prefrontal cortex (vmPFC, area 14), OFC, rostral OFC (rOFC, area 11), pregenual anterior cingulate cortex (pgACC, area 32), posterior cingulate cortex (PCC, area 29), and ventral striatum (VS). We used a previously developed risky choice task with asynchronous presentation of offers (Strait et al., 2014). By presenting offers asynchronously, we were able to characterize neural responses to a single offer in the absence of comparison signals, in the absence of fluctuating attention, which may lead to rapid shifts between the encoded offer (Krajbich et al., 2010; Rich & Wallis, 2016; McGinty et al., 2016; Xie et al., 2018). In all six areas, we found that responses evoked by safe offers were entirely unrelated to responses evoked by equally valued risky ones. Nor did there exist any subpopulation of neurons whose responses showed similar responses to safe and matched value risky offers. In all cases, a simple classifier readily distinguished risky from equally valued safe offers, even in a subpopulation chosen to have minimal firing rate differences. Finally, risky and safe offers occupied dissimilar population subspaces, meaning that risk and safe responses are not just different, they reflect different population organizational frameworks. These results indicate that a basic criterion of abstract value encoding - common responses to equally valued qualitatively different offer types - is violated in all six putative core value regions. The consistency of these results across regions raises the possibility that the brain does not carry abstract value codes anywhere. They therefore support expectancy-based encoding accounts and provide evidence in favor of economic models that eschew abstract value representations (Hayden and Niv, 2021).

## RESULTS

### Behavior

We used the *risky choice task* (Strait et al., 2014; **see Methods**). On each trial, subjects chose between two offers that varied in magnitude and probability (**Figure 1A**). Safe offers (12.5% of offers) provided a small volume of juice (125 *μ*L) with 100% certainty. Risky offers provided either a medium (165 *μ*L, 43.75% of offers) or large (240 *μ*L, 43.75% of offers) volume of juice with a defined probability. The offer types for the two offers were selected independently; for risky offers, the win probability was determined randomly from a continuum of values and indicated unambiguously (0-100%, 1% increments). We collected data from six subjects (*Macaca mulatta*) across a total of 315 sessions comprising 211,884 trials (average 672.6 trials per session).

**Figure 1.**
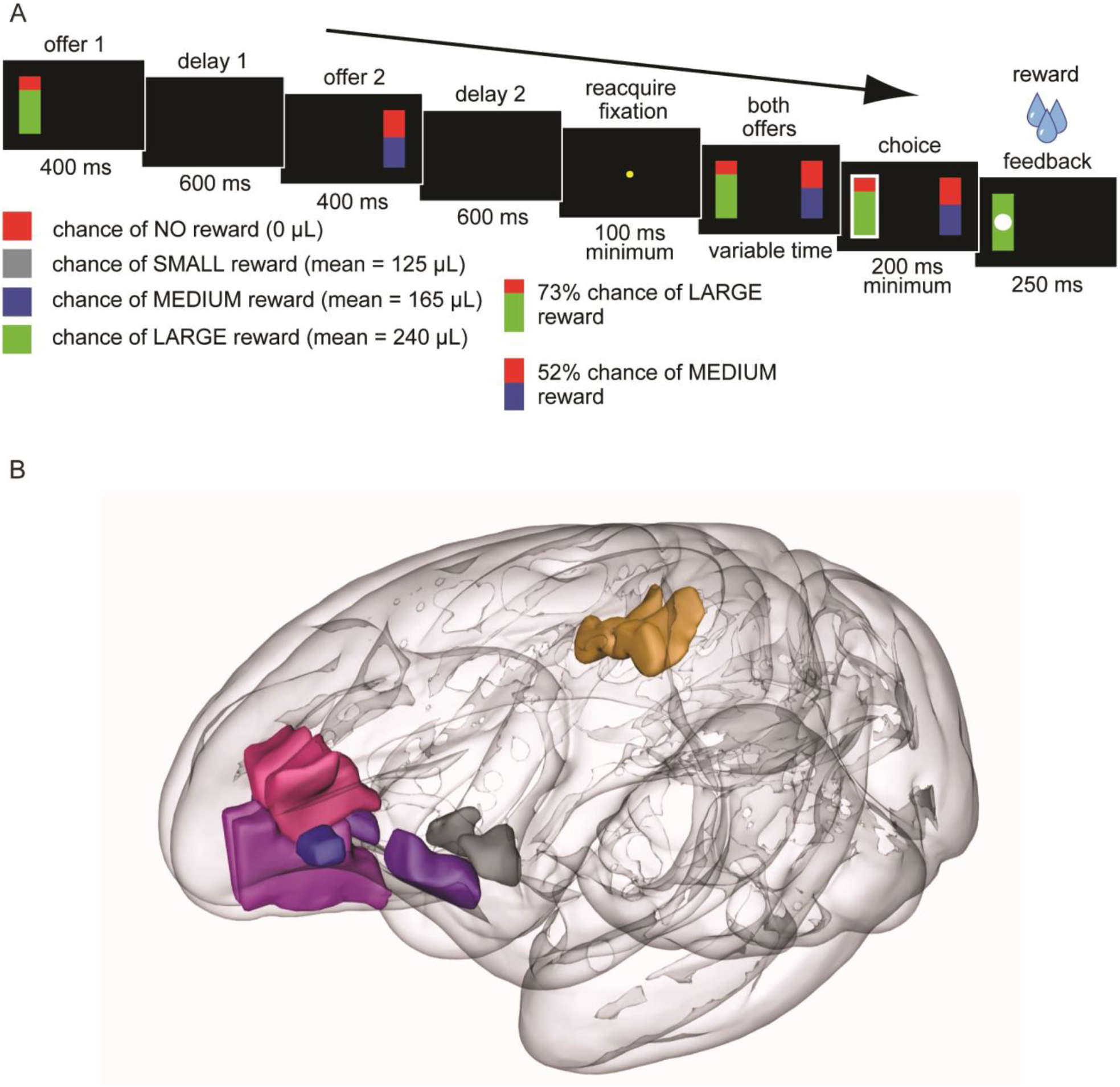
Task, Behavior, and Targeted Structures. **(A)** Structure of our risky choice task (Strait et al., 2014). Each trial begins with a 400 ms presentation of the first offer followed by a 600 ms blank period. Following a 400 ms presentation of the second offer and another 600 ms blank period, a fixation spot appears and, on fixation, both offers appear and the subject selects one by saccade. For each offer, the magnitude of the associated reward (stakes) is indicated by the bottom color (green, high or blue, medium) of the stimulus. The probability of being rewarded is indicated by the size of the green/blue segment. **(B)** Anatomical positions of our brain regions of interest: rOFC (blue), OFC (purple), vmPFC (purple-pink), pgACC (pink), PCC (gold), and VS (grey).

Subjects consistently performed at a high level, were modestly risk-seeking, and did not differ from each other qualitatively (**Figure 2A**). Details of typical behavior in this task are given elsewhere (in greatest detail in Hayden et al., 2010; and in Farashahi et al., 2018 and 2019). Results of these analyses are not repeated here, except to note that subjects’ behavior is quite stable and consistent both within and across sessions (**Figure 2C-D**), and across subjects in this task. Indeed, all tested patterns closely recapitulate those we have observed using this task in the past (ibid.).

**Figure 2.**
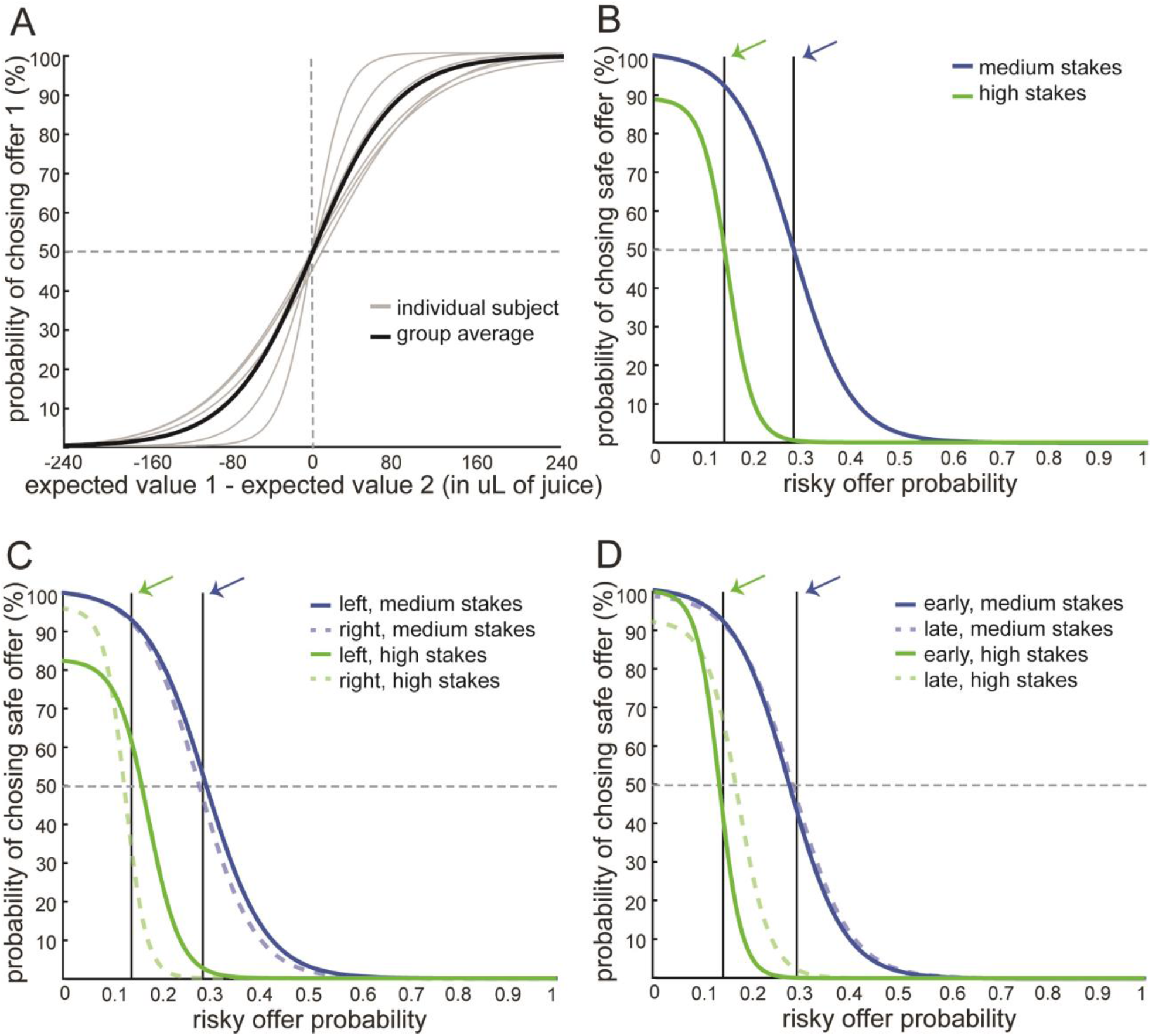
Calculation of equivalent risky and safe values. **(A)** Likelihood of choosing the first offer as a function of its value relative to the second (specifically, for signed value difference). Sigmoid fits of raw binary data shown (see **Methods**). Gray lines: individual subjects; black line: group average. In this and subsequent panels, a horizontal, dashed line indicates the indifference point (the point at which choices are 50/50). **(B)** Likelihood of choosing a safe option as a function of the probability of the risky option for medium (blue) and high (green) stakes offers. All data were analyzed on a subject-by-subject basis, so only data for one example subject (subject B) are shown. Other students showed similar patterns. Vertical black lines (B-D) indicate the probability used as the SV-equivalence point for the subject (the arrow points to the indifference point for medium (blue) and high magnitude (green) risky offers). **(C)** Same as B, except data are separated for left and right offers. Side of presentation does not affect choice much. **(D)** Same as C, except data are separated by trials that were in the first (early) or second (late) half of a session.

The focus of the present study is on comparing responses to safe and risky offers with equivalent subjective values. To identify the relative values of safe offers, we computed the risky-safe indifference point (Hayden et al., 2010). Separately for each subject and separately for medium and large stakes offers, we calculated the likelihood that the subject would choose the safe offer as a function of the probability of the risky offer. We fit the resulting data with a sigmoid curve and calculated the point at which the best-fitting curve crossed the indifference line (**Figure 2**, see **Methods**). We called the value of the risky option that was equivalent to the value of the safe option the indifference point, and assumed that these options were valued equally.

Across all six subjects, the average indifference point for medium magnitude risky offers corresponded to an offer probability of 0.33 +/- 0.05 (standard deviation). A risk-neutral subject would have had an indifference point at 0.76; the fact that the observed value is lower than the optimal indicates that subjects were risk-seeking (Heilbronner & Hayden, 2013; Heilbronner, 2017). Across all six subjects, the average indifference point for high magnitude risky offers corresponded to an offer probability of 0.11 +/- 0.04. A risk-neutral subject would have an indifference point of 0.52). This observation is also consistent with risk-seeking.

As we have observed many times in the past, but never previously published, preferences were strikingly consistent across many contexts. For example, indifference points are similar for the first and second offers (offer 1: medium: 0.34; high: 0.11; offer 2: medium: 0.22; high = 0.11), for offers made early and late in the session (early: medium: 0.29; high: 0.09; late: medium: 0.31; high: 0.13), and when risky offers are appear on the left or right (left: medium: 0.27, high: 0.11; right: medium: 0.36, high: 0.12). Data for an example subject are shown in **Figure 2B-D**.

### Neuronal responses to equally preferred risky and safe offers are unrelated

We recorded responses of 981 neurons in 6 brain regions while our subjects performed the risky choice task: vmPFC (area 14, 156 neurons), OFC (area 13, 157 neurons), rOFC (area 11, 138 neurons), pgACC (area 32, 255 neurons), PCC (area 29/31, 151 neurons), and VS (nucleus accumbens, 124 neurons). Regions are illustrated in **Figure 1B** and anatomical boundaries are provided in the **Methods**. We recorded in two subjects for all areas, although different subjects were used for the different areas (see **Methods**). Detailed analyses of responses to risky offers were reported previously for vmPFC and VS (Strait et al., 2014; Strait et al., 2015). We have not previously examined responses to safe offers.

We reasoned that any neurons that use a common currency code for offer value must produce identical neural responses to subjectively equivalent (that is, equally preferred) risky and safe offers (Padoa-Schioppa & Asaad, 2006; Kennerley et al., 2009; Levy & Glimcher, 2012). Using each individual’s subjective indifference point, we then defined a range of probabilities (+/- 2.5%, total range of 5.0%) and treated all offers within that range as being subjectively equivalent to the safe value. Note that we subsequently checked for robustness by repeating the following analyses using a larger range (+/- 5%, total range of 10%) but because we found no qualitative differences, we do not report those results.

Our analyses focused on the *first offer epoch*, a 500 ms analysis window starting 100 ms after the onset of the first offer. We have used this epoch in all our past research on this and similar tasks (Strait et al., 2014, 2015, and 2016; Azab & Hayden, 2017, 2018, and 2020). We have found that this epoch provides a good characterization of functional responses and allows for fair comparison across brain regions (Strait et al., 2016; Maisson et al., 2020). We used it here for those reasons and because adherence to a single pre-planned epoch of interest reduces the likelihood of inadvertent “p-hacking”.

Neuron vmPFC.70 (**Figure 3A**) showed differing selectivity for safe and risky offers during the first offer epoch. Neuron pgACC.17 (**Figure 3B**), recorded from subject B, showed clear and roughly monotonic tunic for probability of medium and high magnitude risky offers. This means that a downstream decoder could, in principle, readily interpret the firing to identify the probability (and thus, in the context of this task, the value) of a risky offer. It does so because the response of the neuron reflects a single consistent scale for risky offers. However, this scale does not appear to extend to safe offers. Because the safe offer had a subjective value equivalent to 0.28, its mean neural response should have been the same as the response to that offer (0.36 spikes/second) if risky and safe offers used a common scale. Instead, it evoked a mean response of 1.67 spikes/second. (Note that the second quantity is significantly lower than the first, Student’s t-test, t = 4.31, *p* < 0.001). In other words, while this pgACC neuron appears to use a consistent code for risky offers, it does not appear to use the same code for both safe and risky offers. Additional sample cells, from each targeted area, showed positive and negative monotonic tuning while having still clearly responded differently to safe and risky offers (**Figure 3C-F**).

**Figure 3.**
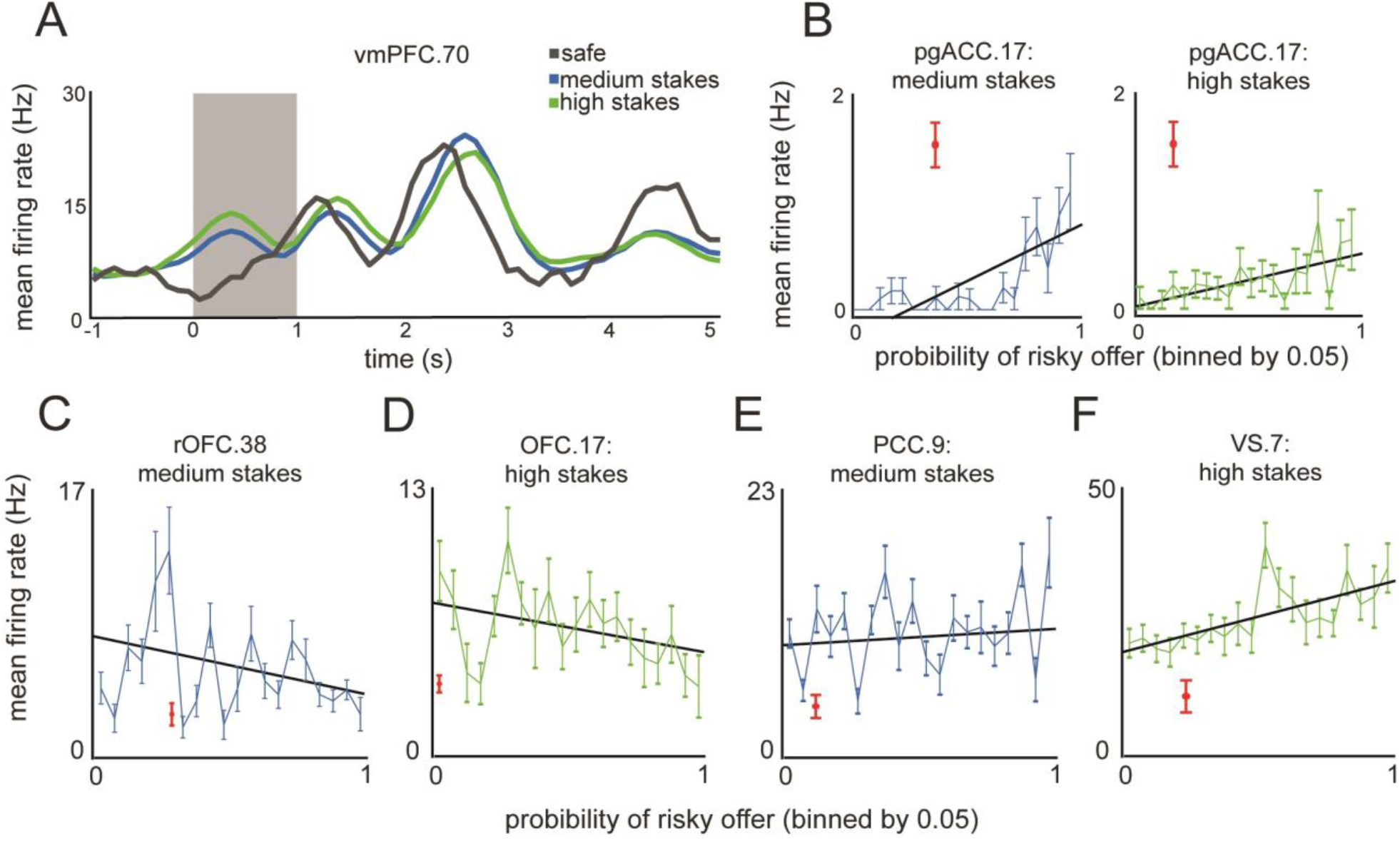
Responses of single neurons. This figures shows the average responses of sample neurons to safe and risky offers of differing values, as well as the average response similarity rates. **(A)** Peristimulus time histogram from mean firing rates of sample neuron vmPFC.70. Each line indicated the average response across offers of a given risk profile (grey: all safe offers; blue: all medium magnitude risky offers; green: all large-magnitude risky offers). The grey shaded box indicates the 1-second period from which the 500-ms epoch 1 analysis window was extracted, where the onset of the first offer is time-locked to zero seconds. **(B)** This is a plot of data collected from a sample neuron in the OFC, which showed a response to safe offers that was statistically different from the response to equivalent risky offers. Depicted are the average responses to medium magnitude (left; blue) and high (right; green) stakes, separated by probability ranges of 0.05. The red point indicates the average response of the given neuron to safe offers (error bars denote the SEM across responses to safe offers). The diagonal black line indicates a fitted regression line, showing positive monotonic tuning. **(C-F)** Same as (B), but demonstrating sample cell responses to an assortment of medium and high stakes offers from across all target areas.

### Responses to safe offers are unrelated to responses to equally valued risky offers

To test whether risky and safe offers are encoded in similar ways in any of our six brain areas, we focused on the key variable, the difference in firing rate response evoked by the two offer types. We call this quantity the *evoked response difference*, or the *delta* for short. If a neuron uses a common currency code for value, then the delta must necessarily be zero. In practice, delta will inevitably deviate from zero due to measurement noise. Specifically, because neuronal responses are stochastic, responses to two different stimuli will necessarily be measured as different even if the true responses are identical. We accounted for this issue by using an analysis approach that combines positive and negative control analyses.

The goal of the **negative control** analysis was to ascertain whether deltas for safe and equally valued risky offers are lower than one would expect by chance. We did this by comparing the difference between the safe response and the response to a randomly chosen probability in the range of 0.0-1.0 (by 0.01 units). In other words, this analysis asks, in effect, whether the safe offer has a special relationship (specifically, a smaller delta) with the equivalently valued risky offer that it does not have with any randomly selected risky offer. If it does not, that implies that risky and safe offers are encoded using unrelated codes.

Consider, for example, neuron vmPFC.10 (subject B). For this neuron, safe offers evoked an average response of 15.71 spikes/second and equivalent medium value risky offers evoked an average response of 19.85 spikes/second. Its delta was therefore the difference between these two numbers, or 4.14 spikes/second. This value is large - roughly 20% of the risky response - but, how large is this value relative to what would be expected by chance? For this neuron, the average delta generated using randomly chosen probabilities (rather than the equivalently valued one) was 2.73 +/- 1.16 spikes/second (this number reflects an average over 1000 randomly sampled probabilities). The observed (true) delta is larger, not smaller, than the control (random) delta, arguing against the hypothesis that this neuron has a common currency code for value. Moreover, these two deltas were not significantly different, (*p* = 0.68, bootstrap test, see **Methods**). In other words, for this neuron, the response evoked by the safe offer is not significantly lower than the value evoked by any random offer - it’s within the range of values one would expect by chance. To put it another way, a downstream decoder would have no way to preferentially associate the safe offer with its equivalently valued risky offer and the evidence suggests the codes for safe and risky offers are entirely unrelated.

We performed this analysis for all recorded neurons in vmPFC. To account for possible scaling differences between responses of different neurons, we used normalized (Z-scored) - firing rates, although the conclusions were unchanged when using raw firing rates. We found that the average normalized delta across neurons for safe and equally preferred medium magnitude risky offers was 0.068 +/- 0.007 (standard deviation) z-score units; the average for the random deltas was similar (0.068 +/- 0.002; these were not different, *p* = 0.487, bootstrap test, see **Methods**). The average normalized delta for safe vs. high magnitude was 0.071; the average for the random deltas was 0.073; these were also not different, *p* = 0.519; **Figure 4B-C**). In other words, neither of these average deltas was significantly smaller than the average deltas between randomly chosen risky offers (using the bootstrapping method; see **Methods**). Note that because there were two conditions, and thus two possibilities of detecting a common currency code, it is appropriate to correct for multiple comparisons; the Bonferroni corrected p-values are *p* = 0.698 and *p* = 0.720, for medium and high magnitude risky offers, respectively. We found the same patterns in the other five structures we examined - most importantly, no area showed a measurable difference between deltas from safe and random risky offers (summarized in **Figure 4B-C**).

**Figure 4.**
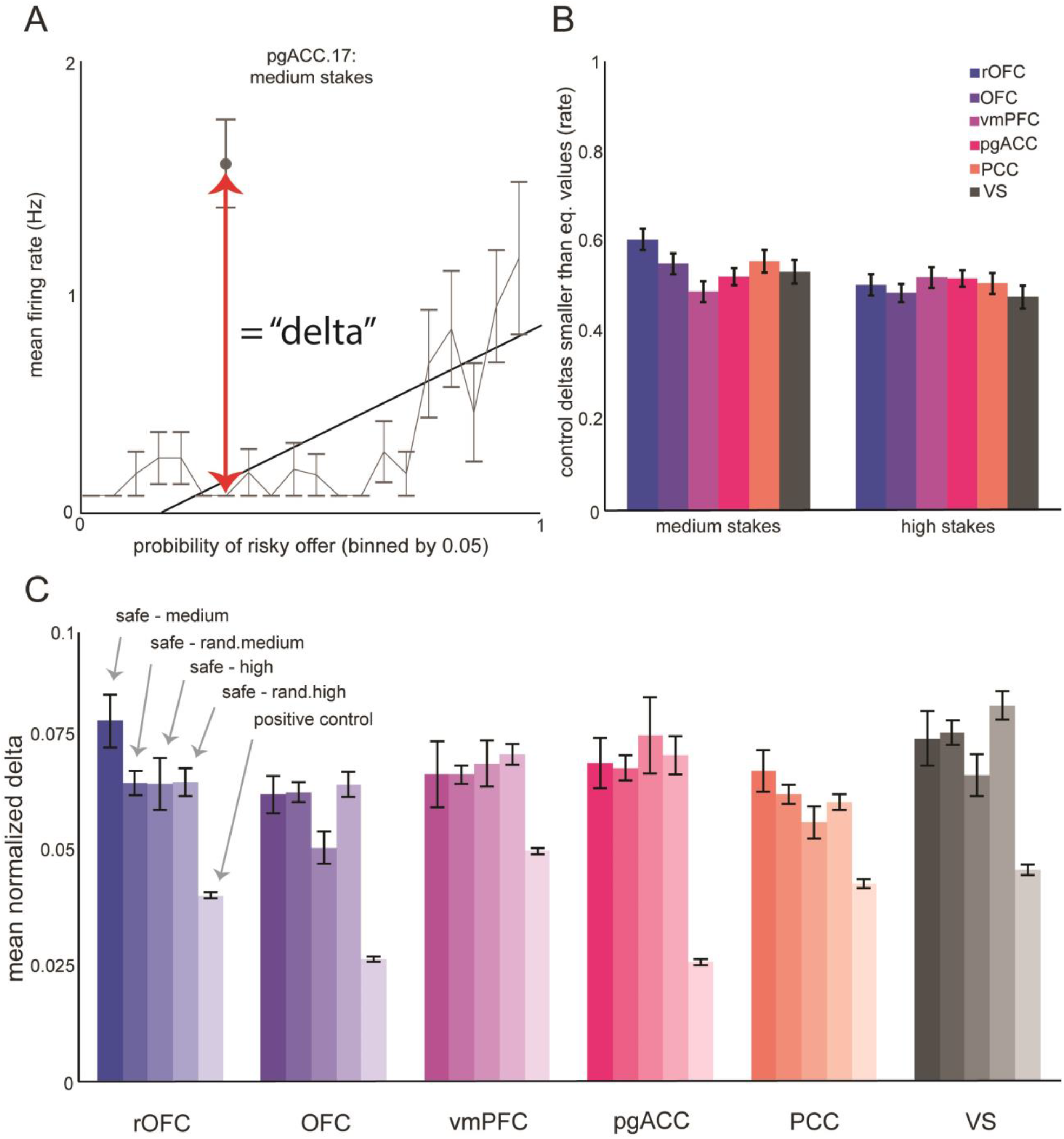
Average risk-safe response difference (delta). **(A)** The same cell depicted in Figure 3B (*left panel*). The red arrow links the responses to the safe and SV-matched risky offers. The absolute value of their difference constitutes the delta. **(B)** The average proportion of deltas, across neurons, between random offers that are smaller than the delta between equivalent safe and risky offers (i.e. the *p-value* from our bootstrap test). The common currency hypothesis would predict that safe - risky deltas are always significantly less than safe - rand.risky deltas. **(C)** The average normalized evoked response difference (delta) for each structure. From left to right (most to least opaque), for each structure: deltas in response to safe v. medium magnitude risky offers, safe v. randomly selected medium offer, safe v. high, safe v. random high, and the positive control deltas. The fact that observed deltas are generally greater than or equal to (never significantly less than) the control deltas (safe - rand.risky) indicates a violation of the common currency hypothesis.

The goal of the **positive control** is to show that our failure to find a difference between the true and random conditions is not itself due to high noise in our sample. In other words, it is theoretically possible that there was enough noise in our measurement of evoked responses that we would have detected a large but spurious delta even if there were no true difference. Our *positive control* analysis is designed to exclude this possibility by confirming we *could* indeed detect a lower than random delta if it existed. (One intuitive way to think about this analysis is that it asks whether we collected sufficiently large numbers of neurons in each area to detect a true common currency code if it existed).

To do this, we randomly assigned trials from SV-matched offers to two sets of equal size, for each neuron. We then computed the delta for each neuron between these two randomized sets. Because these responses are evoked by stochastically identical stimuli, they must have zero true difference. Any measured difference between them gives a measure of what difference we would expect to arise by chance. We reasoned that if the high measured values of our deltas reflected a true violation of common currency coding, they must have necessarily been larger than the responses to the randomly sub-selected sets of risky trials.

We found that the average delta across random sub-selections, computed from vmPFC responses, was 0.051 z-score units. This value was lower than the safe vs. medium magnitude true deltas in vmPFC (which was 0.068, *p* < 0.001, bootstrap test). This positive control delta was also significantly lower than the safe v. high magnitude true delta (0.071, *p* < 0.001, bootstrap test). Similar differences were evident between deltas computed from neural responses in all six structures (*p* < 0.01 in all cases, **Figure 4C**). This analysis indicates that the effects we see are not due to insufficient or noisy data, but instead reflect a true and robust violation of common currency coding.

### No special subpopulation of abstract value encoding neurons

In many cases, common currency coding is not a property of all neurons in a region, but only a subset of specialized abstract value cells (often called offer value cells, e.g. Padoa-Schioppa & Assad, 2006). Our approach, which looks at average response differences in the population, would be sufficient to detect abstract value encoding, even if it were limited to a subpopulation of cells, because those cells would pull down the average delta for the population; our use of the positive control approach (see above) means we can be confident we could detect them even if they were in a very small minority. Thus, that result, while indirect, is still sufficient to cast strong doubt on the idea that there are subpopulations of abstract value cells. Nonetheless, we wanted to more directly test the hypothesis that there is a specialized subpopulation of pure value cells.

We started by reasoning that such a subpopulation, if it exists, will be defined by having an unusually low delta (i.e. low difference between responses evoked by safe offers and equivalently valued risky ones). In theory, that delta would be precisely zero, but (as we state above) because of measurement noise, it will be greater than that; it will nonetheless still necessarily be lower than the delta for the set of non-value neurons. A putative value-coding population, then, can be identified by taking the subset of neurons with the lowest delta value.

Given this logic, we can then use the same approach we developed above for a negative control analysis. Specifically, we can ask whether, given a presumed subpopulation size, the group average delta associated with this subpopulation differs from the group average delta for a matched dummy set of cells identified by using a random probability. If the value is not lower, that argues against the idea that a specialized subset of pure value neurons of a specific set size exists. Because we had no a priori hypotheses about the likely size of such a set, we tested all possible set sizes.

Consider, for example, the possibility that 30% of neurons in OFC (n=47/157) may constitute a unique subpopulation of cells whose responses encode value abstractly. These neurons could be identified by finding those 30% of cells with the lowest difference between responses to safe and equivalent valued risky options (that is, lowest deltas). We then identified these neurons - their average delta was found to be 0.012 z-scored units. Next, we picked a random probability (say, 0.65) and identified another set of 47/157 neurons whose deltas for that probability are minimized. The average random delta for this set of neurons was found to be 0.017 (we can call this the pseudo-delta for 0.65). We next repeated this process 1000 times with 1000 random probabilities, and averaged the resulting pseudo-deltas. The average of these turned out to be 0.015 (**Figure 5A**). Then we asked whether the delta for the equivalent probability was lower than the average pseudo-delta for the random probability (as we would expect if this subpopulation has, or even approximates, an abstract value code). They were not - *p* = 0.171. This result, then, indicates that there is no unique subpopulation of size 30% in OFC.

**Figure 5.**
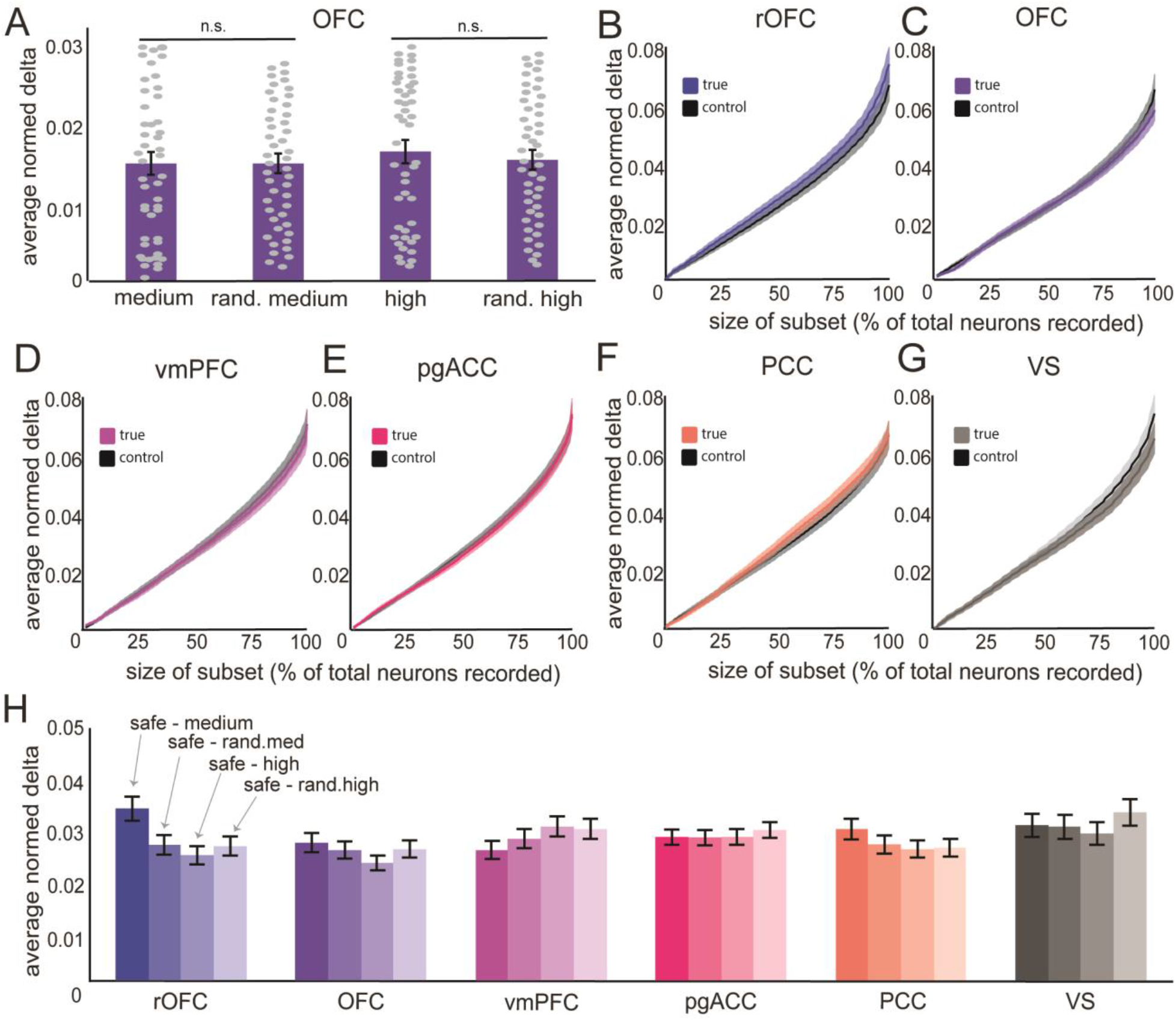
Delta Analysis with best subsets of various sizes. This figure depicts our analysis of response differences in best subsets of neurons. **(A)** For an example structure, OFC, we show the comparison between deltas at example subset size of 30% of recorded OFC neurons. Bars indicate the average normalized deltas for a given comparison: safe v. medium stakes, random medium stakes offers, safe v. high stakes, random high stakes offers. Each dot indicates the delta for a single neuron. Error bars indicate the standard error across the subset. **(B)** Shown is the change in average delta (for visualization only: collapsed across both medium and high stakes comparisons) between safe-risky (blue) and safe-random (black). The lines indicate the average normalized deltas (as in panel A), across subsets from a size of 1-100% of recorded neurons. Shaded ribbons denote the standard error across the subset. *Note again that collapsing across medium and high stakes offers is for visualization only. **(C-G)** same as (C), but for each area of interest. The analysis was performed for safe-medium and safe-high independently. **(H)** For each structure, each bar provides a summary of the average normalized delta across all subset sizes. Error bars represent the standard error across subset sizes.

We next repeated this process for all possible population sizes (1-100%, by 1% increments). (Note that the 100% condition in this test is mathematically equivalent to what we call the negative control delta analysis in the previous section, so this analysis serves as a generalization of that one to all possible set sizes smaller than 100%). As the size of the population increased, naturally so did the average delta across the subpopulation because we necessarily selected more neurons with larger individual deltas (**Figure 5B**). The random deltas also must show the same pattern of increase. The important comparison is to determine whether the safe-risky deltas and safe-random pseudo-deltas, across subsampled population sizes, differ from each other statistically. For each subsampled population size, we computed the deltas both between equivalent and between random offers and performed a Komologorov-Smirnov test, comparing the pair of equivalent and random delta vectors across subset size. We found that there were no significant differences between the equivalent and control deltas as a function of subset size in the other targeted structures (*p* > 0.05 in all cases; **Figure 5C-H**).

### Decodability of safe and equivalently valued risky offers

If the brain specifically encodes value in a common currency manner, then it should not be possible, in principle, to decode safe from risky offers. If, conversely, the brain uses distinct codes for the two categories, then the category (risky vs. safe) should be readily decodable even if the offers are equally valued. We used a standard classifier approach to ask this question. Specifically, we trained a binary support vector machine (SVM) to decode safe from equally valued risky offers based on neural responses (see **Methods**). We then cross-validated the trained model by using it to predict risky vs. safe from responses in a sequestered set.

In OFC, for example, we found that the classifier could readily disambiguate safe from equally valued risky trials (medium magnitude: t = 730.1, *p* < 0.001; high magnitude: t = 780.5, *p* < 0.001). This difference is quite large. For comparison, a t-test comparing 50% to 0% average accuracy along cross-validations, given our pseudo-population method (see **Methods**), and assuming equal variance, would be about the same (t = 701.7). We also observed clear decodability in the other five structures (*p* < 0.001 in all cases; **Figure 6A**).

**Figure 6.**
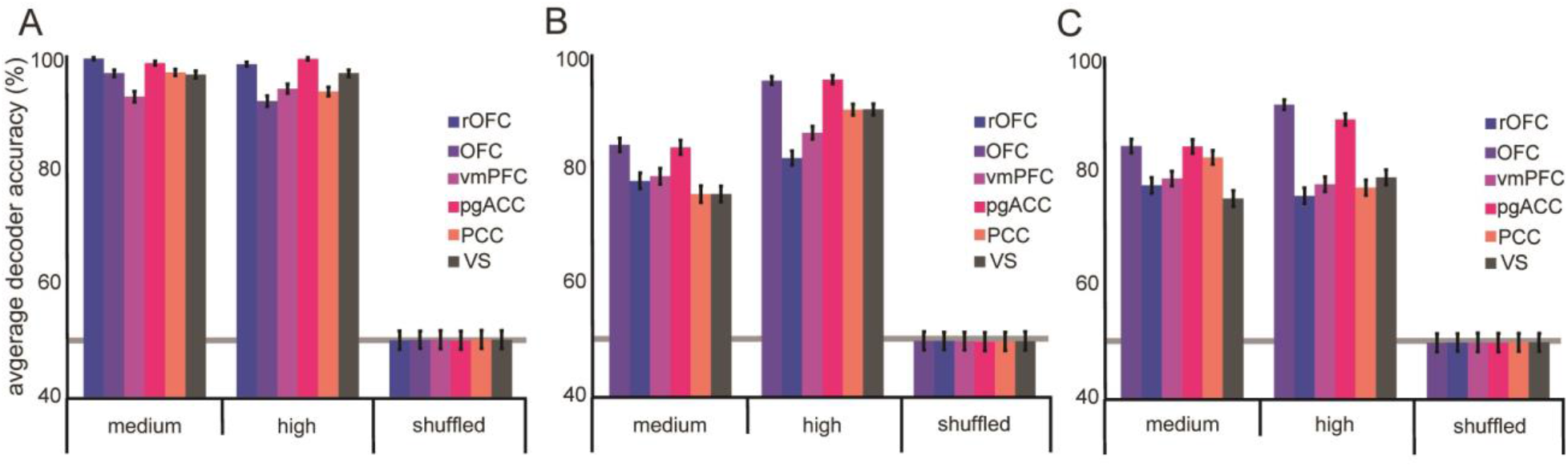
Safe and equivalently valued risky offers are readily decoded. **(A)** Decodability for safe and risky offers is high in all areas for both medium and high stakes gambles. Shuffled data refers to decodability of randomly assigned safe/risky labels to neural responses that are completely shuffled across trials and cells. Error bars indicate the standard error across cross-validations. **(B)** same as A, except that the only neurons used in the decoder were those which, in the first analysis, showed no significant differences in neural responses to safe and equivalent risky offers, and which have differences in responses within the lowest half of the set. **(C)** same as B, except that the only neurons used are those with differences in responses within the lowest quarter of the set

Next, we wanted to confirm that the high decodability rate was not an artifact of using all recorded neurons, including those with stark differences in their responses to equally valued safe and risky offers. We reasoned - as we did above - that there could still be a special subpopulation of pure value cells, such that their abstract encoding of value was obscured by the other neurons. Thus, we identified putative pure-value cells as the set of cells with no significant difference in response to safe and equivalent risky offers (that is, *p* > 0.05). Then, to be even more conservative, we included only the half of these cells with the smallest difference between responses to safe and SV-matched risky offers. In the OFC, this constituted 68/157 neurons (43.3% of recorded cells; rOFC: 60/138 neurons, 43.5%; vmPFC: 66/156 neurons, 42.3%; pgACC: 100/255, 39.2%; PCC: 60/151, 39.7%; VS: 50/124, 40.3%). As before, in OFC we found that the classifier could readily disambiguate the safe from the SV-matched risky trials (medium magnitude: t = 428.5, *p* < 0.001; high magnitude: t = 513.2, *p* < 0.001). This pattern was observed in all five of our structures (*p* < 0.001; **Figure 6B**).

To push even harder against our own conclusions, we conducted the same analysis on an even smaller subset. We included only a quarter of the cells with the smallest difference between responses to safe and SV-matched risky offers. That is, we took the subpopulation that would, by even a very conservative analysis, be most likely to be classified as pure value cells. In the OFC, this constituted 39/157 neurons (24.8% of recorded cells; rOFC: 30/138 neurons, 21.7%; vmPFC: 33/156 neurons, 21.2%; pgACC: 50/255, 19.6%; PCC: 30/151, 19.9%; VS: 25/124, 20.2%). Even so, in OFC, we found that the classifier could readily disambiguate the safe from the SV-matched risky trials (medium magnitude: t = 416.2, *p* < 0.001; high magnitude: t = 391.4, *p* < 0.001). Again, this pattern was still observed in all five of our structures (*p* < 0.001, in all cases; **Figure 6C**).

### Response subspaces for safe and risky offers are different

Responses of ensembles of neurons have correlated variability; these correlations restrict ensemble responses to specific subspaces (Oşan et al., 2007; Gallego et al., 2017). Emerging evidence indicates that neuronal populations can move between subspaces and that subspace reorganization can serve a partitioning function, for example, between motor preparation and execution (Elsayed et al., 2016) or between evaluation and comparison (Yoo & Hayden, 2020). The common currency hypothesis would require the use of a common *abstract value subspace* for different offer types with the same value. Here we asked whether each of our six regions make use of common or distinct subspaces for encoding risky and safe offers.

To do this, we followed an approach to characterize the uniqueness of subspaces for temporally distinct offer epochs, and that is based on methods devised to study subspace reorganization in the motor system (Elsayed et al., 2016; Yoo & Hayen, 2020). We modified that approach to allow for a comparison between safe and risky subspaces within the same epoch. Specifically, we performed a principal component analysis (PCA) on neural responses to both risky and safe offers of equal value. For each subject independently, we projected responses evoked by both safe and equivalent risky offers into the safe offer subspace and computed the total percent variance explained by the top ten principal components. We then averaged the explained variances across subjects (**Figure 7A-B**). We used these projections to quantify the extent to which subspaces were aligned (A_idx_; see **Methods**, Elsayed et al., 2016). This number quantifies the extent of the variance explained in the safe responses, by projecting them into risky response subspace, as a proportion of the variance explained in the safe responses. An alignment index equal to 1.0 would indicate that the variances in risky and safe responses are explained equally well by the principal components that constitute the safe response subspace. That value would suggest common (“aligned”) subspaces and is predicted by the common currency hypothesis. An alignment index equal to zero would indicate that the risky and safe subspaces are strictly orthogonal and an index between 0 and 1 would indicate a partial orthogonality; even partial orthogonality would indicate distinct organizational principles, in violation of the common currency hypotheses.

**Figure 7.**
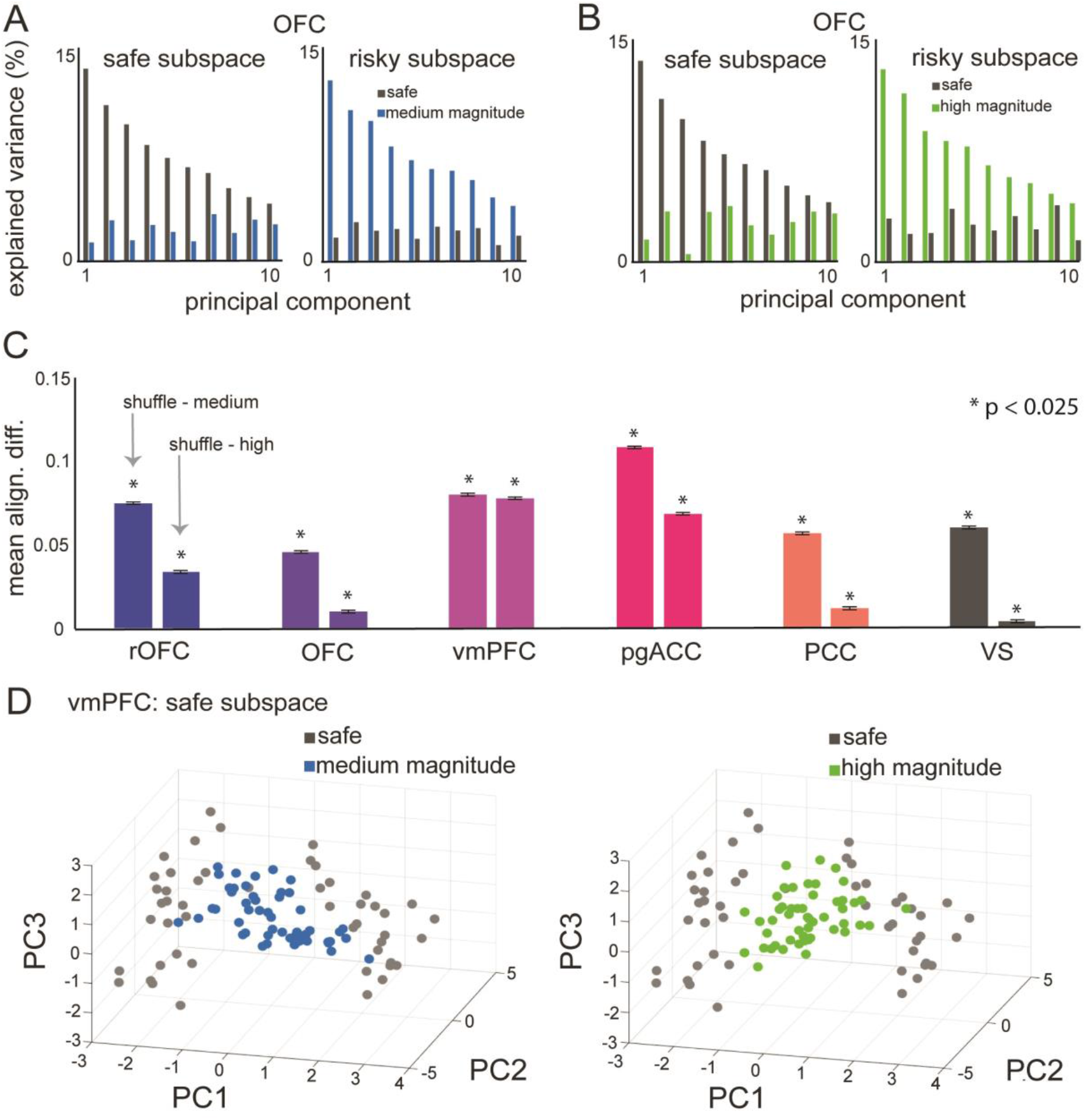
Subspace Alignment between Safe and Equivalent Risky Offers. **(A)** For an example structure, OFC, shows the explained variance (as a proportion of total variance). The left panel shows the explained variance in both safe (grey) and medium stakes (blue) response by projecting both sets of responses into safe response subspace, due to each of the first 10 principal components. The right panel is similar, except that it shows explained variance by projecting both sets of responses into risky response subspace. **(B)** same as (A), except that it shows high stakes (green) offer projections into safe subspace (left) and safe (grey) offer projections into high stakes subspace (right). **(C)** a summary of the difference between the average shuffle alignment index and either the safe-medium (left bar) or the safe-high (right bar) alignment index. Error bars indicate the standard error across computed differences. **(D)** For an example structure, OFC, *left panel*: projections of medium stakes (blue) responses and safe (grey) responses, into safe subspace. *Right panel*: projections of high stakes (green) responses and safe (grey) responses, into safe subspace.

We found that OFC safe and medium magnitude risky response subspaces had an alignment index of A_idx_ = 0.21. Safe and high magnitude subspaces had an alignment of A_idx_ = 0.25. The critical question is whether this value is significantly less than 1.0, which would indicate at least partial orthogonalization. As a control, to determine the significance of our results, we reasoned that breaking the structure of the within-time covariance, while maintaining the within-neuron covariance, should produce subsets that are more orthogonal than would be expected by the within-neuron covariance structure alone. That is, any alignment index at or below this threshold would be considered primarily orthogonal. To do this, we shuffled the data from across all three response matrices (safe, matched-medium, and matched-large) and computed the alignment index between shuffled sets (see **Methods**). We repeated this process over 1000 iterations. Then, to test for significance, we computed the 95% confidence interval across iterations; a value outside of this range can therefore be said to be significant at p < 0.025 (two-tailed t-test). We found that the average shuffled alignment index in OFC was A_idx_ = 0.276. Both the safe-medium and safe-high alignment indexes were below the 95% confidence interval (0.274 - 0.278). In other words, response subspaces for safe and equally valued risky offers in OFC are more orthogonal than we would expect by chance given the inherent statistical properties of our dataset (**Figure 7C**). We found similar results in all structures (safe-medium and safe-high was below the 95% confidence interval, in all cases; p < 0.025).

Our results suggest that the subspaces are generally more orthogonal than they are aligned. However, we wanted to confirm that regardless of the degree of alignment, the safe, medium, and high stakes neural responses are readily separable when projected into the safe offer subspace. To do this, we again projected each of the neural response matrices into the safe offer subspace (**Figure 7D**). We then performed a multinomial logistic regression (to discriminate class based on linear trends within the projections), using the projections onto the top 10 principal components to predict whether the offer was safe, medium, or high stakes. We found that, in all structures, the regression model was significantly able to discriminate between offer types (rOFC: F = 13.36, *p* < 0.001; OFC: F = 13.33, *p* < 0.001; vmPFC: F = 13.32, *p* < 0.001; pgACC: F = 13.26, *p* < 0.001; PCC: F = 13.26, *p* < 0.001; VS: F =13.29, *p* < 0.001). Since the factors used in the regression model are all projections into safe subspace, these results suggest that safe and risky offers operate in highly separable hyperplanes. That is, safe and risky information can likely be easily demixed in a given subspace, in striking violation of the common currency hypothesis.

## DISCUSSION

We examined responses of neurons in six core reward regions to risky and safe offers. By using a large number of risky offers for each neuron, we were able to identify, post-hoc, a subset of risky offers with values equivalent to those of the safe offers. We were then able to ask whether responses to safe and subjective value-matched risky offers were the same, as predicted by common currency models. They were not. Indeed, responses elicited by safe offers were no more similar to those elicited by equally valued risky offers than to any other risky offer in our offer set, indicating that the neural codes used for risky and safe options are unrelated. Moreover, a simple classifier could easily distinguish safe from risky options, even when limited to the subset of neurons most likely to carry a common currency code. Finally, safe and risky options elicited responses in distinct non-collinear subspaces. Together these results provide strong evidence against the idea that the brain makes use of a common currency code for risky and safe offers.

Even among those who favor the common currency hypothesis, there is some debate about the most likely region or regions in which common currency coding is likely to occur (Rangel et al., 2008; Kable & Glimcher, 2009; Padoa-Schioppa, 2011; Rushworth et al., 2011; Wunderlich et al., 2012). One tradition focuses on the OFC and most research in that tradition focuses on area 13 (Tremblay & Schultz, 1999; Wallis, 2007; Padoa-Schioppa, 2011). Some work also implicates area 11 (Rudebeck & Murray, 2014). We tested both areas. Another tradition, much of it based on human neuroimaging, favors vmPFC, which is quite different from OFC, both in terms of connectivity and other possible functions (Blair, 2008; Noonan et al., 2011; FitzGerald et al., 2012; reviewed in Levy & Glimcher, 2012, Bartra et al., 2013, and in Clithero & Rangel, 2014). The primate homologue of vmPFC is unclear - it may be area 14 or it may be area 32 (Myers-Schulz & Koenigs, 2012; Neubert et al., 2015). We tested both. A third tradition emphasizes the likely importance of PCC for common currency functions (McCoy & Platt, 2005; Kable & Glimcher, 2007). Finally, another major theory focuses on the ventral striatum, especially on the NAc core region (Beck et al., 2009; Knutson et al., 2009; Staudinger et al., 2009; Cai et al., 2011; Strait et al., 2015). Our study, which uses the same task in all regions, and finds the same lack of common currency coding, therefore represents a relatively complete list of putative common currency regions, and raises the possibility that such a code does not exist anywhere in the brain. Note, however, that our results do not demonstrate that these areas have identical functions; instead they indicate that whatever value signals are available in these regions are not amodal in our risky choice task.

The idea of a common currency representation argues that there is a single final common pathway for value that can feed into - but is conceptually distinct from - action selection. If the brain does not make use of a common currency, then how can we compare values of dissimilar things? There are many possible answers (Vlaev et al., 2011; Hayden & Niv, 2020). Process models of choice that eschew value representations include heuristic approaches, sampling-based approaches, and embodied/premotor theories. They also include, for example, distributed choice implementations such as that of bee and ant swarms (Seeley et al., 1991; Marshall et al., 2009; Seeley et al., 2012; Bose et al., 2017; Pirrone et al., 2018). In bee swarms, no single bee has access to the value of an option on a universal scale; instead the comparison is made in an indirect manner (Seeley et al., 1991; Seeley et al., 2012). What these approaches have in common is that they do not involve direct comparison of values; instead, they achieve choice by indirect manners. Such approaches tend to be well tailored to natural decision-making contexts in which single options often appear, and the decision must occur without knowledge of the values of the alternatives (Hayden, 2017). In any case, common currency is one of several possible ways to describe the patterns of choice observed in human and non-human animal decision-makers.

Previous results have demonstrated amply that neurons in the brain have responses that correlate with the values of offers that differ in the ratio of their components (e.g. Padoa-Schioppa & Assad, 2006). For example, we have shown that two matched risky options, high-probability low stakes and low-probability high stakes, if equally valued, produced responses along a value axis in vmPFC and VS (Strait et al., 2014; Strait et al., 2015). Such responses satisfy one prediction of the common currency hypothesis. Our new results presented here do not vitiate these earlier ones; instead, they demonstrate the limitation of past findings - that they did not test enough conditions to fully falsify the common currency theory. Specifically, our current results suggest that a second criterion for common currency coding, a common code for options that differ in kind, not just in ratio, is not satisfied in any of the major proposed reward regions. These results, then, suggest that key reward areas can integrate across dimensions using a single scale but may use different scales for qualitatively different offer types. Note that, in a trivial sense, risky and safe options are different ratios of risk and stakes - but it is clear they differ psychologically. For example, the risky option affords an opportunity for learning/adjustment, while the safe does not. The risky option may trigger different processes, such as anxiety or savoring (Lopes, 1987; Loewenstein et al., 2001).

Perhaps the most intriguing result is our finding that different equally valued stimuli are encoded in non-collinear response subspaces. The brain can make use of response subspaces to keep pieces of information separate. For example, premotor plans can be kept in an output-null subspace so that motor planning can occur without risking triggering a premature action (Kaufman et al., 2014; Elsayed et al., 2016). We have recently explored the idea that subspace orthogonalization in reward regions may also be used to sequester evaluation from comparison in time (Yoo and Hayden, 2020). Our results suggest that subspace orthogonalization may have a second benefit for reward regions - in particular, that it can segregate qualitatively different option types (Semedo et al., 2019; Stokes et al., 2020). That in turn may serve a categorization function - that is, it may help the brain identify, to downstream decoders, which stimulus type it is encoding. The decoder, then, would have the ability to know whether the stimulus presented was risky or safe, and which decoding procedure to use. This idea, however, is speculative, and further research will be required to test it.

Our work relates to a major ongoing debate about the function of OFC, a set of regions whose functions have long been discussed (reviewed in Gardner & Schoenbaum, 2020). One prominent theory links it to value-specific functions, most notably in representing the values of options (Wallis, 2007; Padoa-Schioppa, 2011; Rudebeck & Murray, 2014). However, another theory holds that it serves primarily to implement expectancy signalling - that is, it encodes the properties of potential outcomes associated with reward (reviewed in Gardner & Schoenbaum, 2020). These outcomes may include the reward value but would include all other features. For this reason, qualitatively different offers that have the same value should, according to the expectancy theory, produce unrelated neural responses (Wang & Hayden, 2017). The expectancy theory, more generally, is associated with the idea that the function of OFC is to implement a cognitive map of task space (Wilson et al., 2014; Shuck et al., 2016; Behrens et al., 2018; Schuck & Niv, 2019). Our results, then, not only endorse the expectancy and cognitive map theories of OFC function, but suggest they may apply to other putative reward regions as well.

## Acknowledgements

We thank Meghan Castagno Pesce, Marc Mancarella, Caleb Strait and Tommy Blanchard for assistance with data collection, Sarah Heilbronner for help with anatomy, and the rest of the Hayden and Zimmermann labs for valuable discussions. This research was supported by a National Institute on Drug Abuse grant P30 DA048742-01A1 (to BYH and JZ), a National Institute for Biomedical Imaging Grant P41 EB027061 (to BYH and JZ), and a UMN AIRP award (to BYH and JZ).

## METHODS

### Surgical procedures

All procedures were approved by either the University Committee on Animal Resources at the University of Rochester or the IACUC at the University of Minnesota. Animal procedures were also designed and conducted in compliance with the Public Health Service’s *Guide for the Care and Use of Animals*. Six male rhesus macaques (*Macaca mulatta*) served as subjects. A small prosthesis head fixation was used. Animals were habituated to laboratory conditions and then trained to perform oculomotor tasks for liquid rewards. We place a Cilux recording chamber (Crist Instruments) over the area of interest (see *Behavioral tasks* for breakdown). We verified positioning by magnetic resonance imaging with the aid of a Brainsight system (Rogue Research). Animals received appropriate analgesics and antibiotics after all procedures. Throughout both behavioral and physiological recording sessions, we kept the chamber with regular antibiotic washes and we sealed them with sterile caps.

### Recording sites

We approached our brain regions through standard recording grids (Crist Instruments) guided by a micromanipulator (NAN Instruments). All recording sites were selected based on the boundaries given in the Paxinos atlas (Paxinos et al., 2008). In all cases we sampled evenly across the regions. Neuronal recordings in OFC were collected from *subjects P and S*; recordings in rOFC were collected from *subjects V and P*; recordings in vmPFC were collected from *subjects B* and *H*; recordings in pgACC were collected from *subject B and V*; recordings from PCC were collected from *subject P and S*; and recording in VS were collected from *subject B and C*. Specifically (see **Figure 1B**):

We defined **rOFC 11** as lying within the coronal planes situated between 34.05 and 42.15 mm rostral to the interaural plane, the horizontal planes situated between 4.5 and 9.5 mm from the brain’s ventral surface, and the sagittal planes between 3 and 14 mm from the medial wall. The coordinates correspond to area 11 in Paxinos et al. (2008).

We defined **OFC 13** as lying within the coronal planes situated between 28.65 and 34.05 mm rostral to the interaural plane, the horizontal planes situated between 3 and 6.5 mm from the brain’s ventral surface, and the sagittal planes between 5 and 14 mm from the medial wall. The coordinates correspond to area 13m in Paxinos et al. (2008).

We defined **vmPFC 14** as lying within the coronal planes situated between 29 and 44 mm rostral to the interaural plane, the horizontal planes situated between 0 and 9 mm from the brain’s ventral surface, and the sagittal planes between 0 and 8 mm from the medial wall. These coordinates correspond to area 14m in Paxinos et al. (2008).

We defined **pgACC 32** as lying with the coronal planes situated between 30.90 and 40.10 mm rostral to the interaural plane, the horizontal planes situated between 7.30 and 15.50 mm from the brain’s dorsal surface, and the sagittal planes between 0 and 4.5 mm from the medial wall (**Figure 1B**). Our recordings were made from central regions within these zones, which correspond to area 32 in Paxinos et al. (2008).

We defined **PCC 29/31** as lying within the coronal planes situated between 2.88 mm caudal and 15.6 mm rostral to the interaural plane, the horizontal planes situated between 16.5 and 22.5 mm from the brain’s dorsal surface, and the sagittal planes between 0 and 6 mm from the medial wall. The coordinates correspond to area 29/31 in Paxinos et al. (2008).

We defined **VS** as lying within the coronal planes situated between 20.66 and 28.02 mm rostral to the interaural plane, the horizontal planes situated between 0 and 8.01 mm from the ventral surface of the striatum, and the sagittal planes between 0 and 8.69 mm from the medial wall. Note that our recording sites were targeted towards the nucleus accumbens core region of the VS.

We confirmed recording location before each recording session using our Brainsight system with structural magnetic resonance images taken before the experiment. Neuroimaging was performed at the Rochester Center for Brain Imaging on a Siemens 3T MAGNETOM Trio Tim using 0.5 mm voxels. We confirmed recording locations by listening for characteristic sounds of white and gray matter during recording, which in all cases matched the loci indicated by the Brainsight system with an error of ∼1 mm in the horizontal plane and ∼2 mm in the z-direction.

### Electrophysiological techniques

Either single (FHC) or multi-contact electrodes (V-Probe, Plexon) were lowered using a microdrive (NAN Instruments) until waveforms between one and three neuron(s) were isolated. Individual action potentials were isolated on a Plexon system (Plexon, Dallas, TX) or Ripple Neuro (Salt Lake City, UT). Neurons were selected for study solely on the basis of the quality of isolation; we never preselected based on task-related response properties. All collected neurons for which we managed to obtain at least 300 trials were analyzed; no neurons that surpassed our isolation criteria were excluded from analysis.

### Eye-tracking and reward delivery

Eye position was sampled at 1,000 Hz by an infrared eye-monitoring camera system (SR Research). Stimuli were controlled by a computer running Matlab (Mathworks) with Psychtoolbox and Eyelink Toolbox. Visual stimuli were colored rectangles on a computer monitor placed 57 cm from the animal and centered on its eyes (Fig. 1*A*). A standard solenoid valve controlled the duration of juice delivery. Solenoid calibration was performed daily.

### Behavioral tasks

Six monkeys performed in the risky choice task. Both tasks made use of vertical rectangles indicating reward amount and probability. We have shown in a variety of contexts that this method provides reliable communication of abstract concepts such as reward, probability, delay, and rule to monkeys (Blanchard et al., 2015; Sleezer et al., 2016; Mehta et al., 2019).

### Risky choice task

The task presented two offers on each trial. A rectangle 300 pixels tall and 80 pixels wide represented each offer (11.35° of visual angle tall and 4.08° of visual angle wide; Fig. 2*A*). Two parameters defined gamble offers, *stakes* and *probability*. Each gamble rectangle was divided into two portions, one red and the other either gray, blue, or green. The size of the color portions signified the probability of winning a small (125 μl, gray), medium (165 μl, blue), or large reward (240 μl, green), respectively. We used a uniform distribution between 0 and 100% for probabilities. The size of the red portion indicated the probability of no reward. Offer types were selected at random with a 43.75% probability of blue (medium magnitude) gamble, a 43.75% probability of green (high magnitude) gambles, and a 12.5% probability of gray options (safe offers).

On each trial, one offer appeared on the left side of the screen and the other appeared on the right. We randomized the sides of the first and second offer. Both offers appeared for 400 ms and were followed by a 600-ms blank period. After the offers were presented separately, a central fixation spot appeared and the monkey fixated on it for 100 ms. Following this, both offers appeared simultaneously and the animal indicated its choice by shifting gaze to its preferred offer and maintaining fixation on it for 200 ms. Failure to maintain gaze for 200 ms did not lead to the end of the trial but instead returned the monkey to a choice state; thus monkeys were free to change their mind if they did so within 200 ms (although in our observations, they seldom did so). Following a successful 200-ms fixation, the gamble was resolved and the reward was delivered. We defined trials that took > 7 sec as inattentive trials and we did not include them in the analyses (this removed ∼1% of trials). Outcomes that yielded rewards were accompanied by a visual cue: a white circle in the center of the chosen offer. All trials were followed by an 800-ms intertrial interval with a blank screen.

### Estimation of subjective value equivalence

We calculated the indifference point between safe and risky offers. For each subject, independently, we fitted a sigmoidal function to the distribution of choices (safe or risky) across the full range of risky offer probabilities.

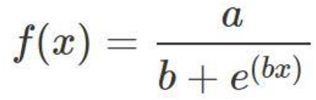

where x = the probability associated with the risky offer, and *f*(x) = the likelihood of choosing the safe offer; a (the maximum value of the curve) and b (the growth rate; steepness) are coefficients of the function, estimated by the fitting procedure for maximizing *R*^*2*^. Using the fitted sigmoidal function, we then estimated the value of x needed to produce a safe choice likelihood of 0.5; that is, a risky offer choice is equally likely as a safe choice. This is called the *indifference point*, and it allows us to calculate the risky offer value that is considered by the subject to be of equivalent value to that of the safe offer. We performed this analysis separately for medium and high stakes gambles, and separately for each subject.

### Statistical methods

We constructed peristimulus time histograms by aligning spike rasters to the presentation of the first offer and averaging firing rates across multiple trials. We calculated firing rates in 20-ms bins but we analyzed them in longer (500 ms) epochs. Some statistical tests of neuronal activity were only appropriate when applied to single neurons because of variations in response properties across the population.

### Pure value cell analysis

For each neuron, we separated the firing rates in response to offer 1 by whether they corresponded to a safe or risky offer. Each epoch consisted of a 500 ms window, beginning 100 ms after the onset of the corresponding offer, which we have found in previous work to roughly correspond with the stimulus-response lag time (Strait et al., 2014).We calculated the mean firing rates for each neuron across all safe or risky trials. We then performed a t-test between firing rate vectors in response to safe and risky offers with equivalent value. We calculated the remaining proportion of total neurons in a given structure after neglecting those with significantly different firing rates in response to equal offers. We then repeated this analysis for firing rates in response to safe offers and a randomly chosen risky offer. We repeated this control step 1000 times. The proportion of sample neurons remaining after accounting for those with statistically differing responses constituted the *true positive* rate, while the average proportion across the 1000 random samples constituted the *false positive* rate, or the rate at which we should expect to see statistically similar firing rates purely by chance. To confirm statistical significance, we determined the proportion of bootstrapped *false positive* rates that were less than the *true positive* rate. A statistically significant *true positive* rate would be larger than at least 950 of the 1000 bootstrapped samples. Therefore, the *p-value* is equal to the proportion of *false positive* rates that are larger than *true positive* rates (or, the difference between one and that the number of *false positive* rates that are smaller than the *true positive* rate). That is, the larger the *true positive* rate is relative to the 1000 samples of *false positive* rates, the smaller the *p-value* will get.

### Response difference (delta) analysis

For each neuron, we separated the firing rates in response to offer 1 by whether they corresponded to a safe or risky offer. We calculated the mean firing rates for each neuron across all trials safe and risky trials. We then computed the absolute value of the difference between mean firing rates in response to safe and equivalent risky offers. To determine significance, we performed a 1000-sample bootstrap of randomly selected risky offers, computed the delta, ranked the deltas in ascending order and asked how many of these 1000 control deltas were less than the true delta. We next defined subsets of neurons ranging from 1-100% of all recorded neurons in a structure. We performed these same calculations, and ordered the differences from least to greatest. For each subset, we took the corresponding number of neurons with the smallest response difference (i.e. the best subset) and compared them to a control population (calculated the same was as described previously). We then performed a Komolgorov-Smirnov test to compare the average response difference to equal offers across all subset sizes with the response difference to random offers.

### Decoding analysis

We built a pseudo-population of pseudo-trials. First, for each epoch, we isolated firing rate responses to the safe offers and the equivalent risky offers. Then, we collapsed the firing rates for each trial into an average for the 500 ms period. We randomly selected 1000 samples for each neuron, under both risk conditions, resulting in two *n* X 1000 matrices (one for each label level), where n represented the number of neurons recorded from each region. This constituted the pseudo-population of pseudo-trials. To execute the decoder, each matrix was split in half and concatenated with the half from the other label. We used one of these matrices to train a binary support vector machine, the other was used for cross-validation. We used the trained model to predict the binary label (safe or risky) for each pseudo-trial in the cross-validation set. We then compared the predicted label to the known label and an accuracy rate was calculated across predictions. This process was repeated 1000 times for each structure and epoch to get a distribution of accuracy rates. Thus, the standard error of the mean, used in displaying the error bars, represents the standard error over the variance of the cross-validations. Additionally, the exact process was repeated on randomly shuffled data, to confirm that expected prediction accuracy was 50% when randomized.

### Subspace alignment

We followed the procedure described in Yoo and Hayden (2020). Specifically, for each structure, we separated offers and neural responses by their risk profile (safe and equivalent risky offer of medium and high magnitudes), as described previously. For each neuron, we identified two factors to incorporate into a single condition: time and offer position. Time included the same 500 ms period following the onset of offer 1 and preceding the onset of offer 2. Time was segmented into 20 ms bins. For each 20 ms bin, we computed the mean firing rate across trials on which the offer was positioned on either the left or right of the screen. Thus, we constructed a condition (time X offer position) X neuron matrix of mean firing rates; that is, a 50 X n-neurons matrix. One such matrix was constructed for safe offers, one for medium, and one for high magnitude risky offers of equivalent value. Firing rates in prefrontal areas of macaques tend to be sparse. So, we smoothed these matrices using a gaussian filter, with a sigma equal to one. We then normalized the smoothed matrices, by computing the z-score within each cell, to account for differences in encoding scaling between neurons.

Next, we performed a principal component analysis, using eigenvalue decomposition, on the safe response matrix, providing a transformation matrix into which we projected both the safe response matrix and each of the risky response matrices. We computed the explained variance due to each of the principal components. We performed the same process of dimensionality reduction for each of the risky offers, projecting both the safe response and corresponding risky response data into the resulting principal component spaces (medium magnitude and high magnitude risky response each into their own principal component spaces). To determine if the subspaces were aligned, we computed an alignment index:

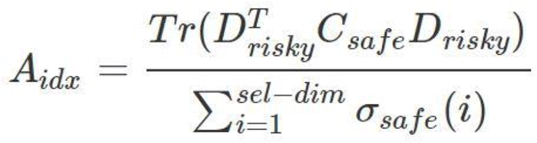

where Tr() is the sum along the diagonal entry, sel-dim = the number of selected principal components (or ten, in the current study), D_risky_ are the set of top sel-dim eigenvectors, C_safe_ is the covariance matrix for the safe responses, σ_safe_(i) is i-th singular value of C_safe_. Essentially, the variance explained in safe response by the top ten principal components of the risky responses is normalized against the sum of the variance explained by the top ten principal components of the safe responses. Note that we also performed this calculation using both the top 4 and top 7 principal components. This control did not change the results of the significance tests, and so they are not reported.

To determine the significance of the alignment index we performed a shuffle procedure. We assume, in this procedure, that neural responses adhere to a fixed correlation structure (Elsayed et al., 2016). Thus, we tested whether the safe and risky subspaces were more or less orthogonal, relative to randomly sampling within the space of this fixed correlation structure. We concatenated all safe and risky offer data into a single matrix. We then computed the covariance matrix, and performed an eigenvalue decomposition for the covariance matrix. We then randomly sampled subspaces that were aligned to the fixed correlation structure of the response space, using a method described by Elsayed et al. (2016), as follows:

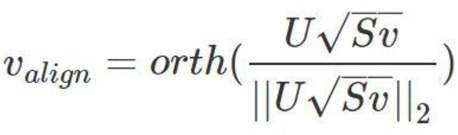

where U and S are the eigenvectors and eigenvalue matrices, respectively, of the computed covariance matrix. A matrix (v) was drawn from a normal distribution with a mean = 0.0 and variance = 1.0. *Orth*() computes the orthonormal basis of the projected matrix. This process essentially maintains the neuronal covariance structure of the original covariance matrix used for the eigenvalue decomposition. We repeated this process across 1000 iterations and computed the alignment index for each, according to the above description. We then calculated the average alignment index and the 95% confidence intervals across the 1000 iterations.

### Multinomial Logistic Regression

To follow-up on the alignment index analysis, we used the projections of safe, medium, and high stakes risky offers into safe subspace. We selected the top 10 results principal components, consistent with the alignment index and constituting no less than 80% of the total explained variance. We then built a multinomial logistic regression model, in which projections onto the first 10 principal components, from of the offer types, were used as simultaneous predictors. The model then set these factors as predictors of a categorical class variable, labeling each offer type. The resulting model was then tested for significance using the standard F-statistic for regressions, against an alpha of 0.05.

## References

Azab, H. and Hayden, B. Y. (2017). Correlates of decisional dynamics in the dorsal anterior cingulate cortex. PLoS Biology. https://doi.org/10.1371/journal.pbio.2003091

Azab, H. and Hayden, B. Y. (2018). Correlates of economic decisions in the dorsal and subgenual anterior cingulate cortices. European Journal of Neuroscience. https://doi.org/10.1111/ejn.13865

Azab, H. and Hayden, B. Y. (2020). Partial integration of the components of value in anterior cingulate cortex. Behavioral Neuroscience, 134, 296–308.

Barseghyan, L., Molinari, F., O’Donoghue, T., and Teitelbaum, J. C. (2013). The nature of risk preferences: Evidence from insurance choices. American Economic Review, 103.

Bartra, O., McGuire, J. T., Kable, J. W. (2013). The valuation system: A coordinate-based meta-analysis of BOLD fMRI experiments examining neural correlates of subjective value. NeuroImage, 76, 412–427.

Beck, A., Schalgenhauf, F., Wüstenberg, T., Hein, J., Kienast, T., … and Wrase, J. (2009). Ventral striatal activation during reward anticipation correlates with impulsivity in alcoholics. Biological Psychiatry, 66, 734–742.

Behrens, T. E. J., Muller, T. H., Wittington, J. C. R., Mark, S. Baram, A. B. … and Kurth-Nelson, Z. (2018). What is a cognitive map? Organizing knowledge for flexible behavior. Neuron, 100, 490–509.

Blair, R. J. R. (2008). The amygdala and ventromedial prefrontal cortex: functional contributions and dysfunction in psychopathy. Phil. Trans. R. Soc. B, 363, 2557–2565

Blanchard, T. C., Hayden, B. Y., and Bromber-Martin, E. S. (2015). Orbitofrontal cortex uses distinct codes for different choice attributes in decisions motivated by curiosity. Neuron, 85, 602–614.

Bose, T., Reina, A., and Marshall, J. A. R. (2017). Collective decision-making. Current Opinion in Behavioral Sciences, 16.

Cai, X., Kim, S., and Lee, D. (2011). Heterogeneous coding of temporally discounted values in the dorsal and ventral striatum during intertemporal choice. Neuron, 69, 170–192.

Clithero, J. A. and Rangel, A. (2014). Informatic parcellation of the network involved in the computation of subjective value. Social cognitive and affective neuroscience, 9, 1289–1302.

Dorris, M. C. and Glimcher, P. W. (2004). Activity in posterior parietal cortex is correlated with the relative subjective desirability of action. Neuron, 44.

Elsayed, G. F., Lara, A. H., Kaufman, M. T., Churchland, M. M., and Cunningham, J. P. (2016). Reorganization between preparatory and movement population responses in motor cortex. Nature Communications, 7.

Farashahi, S., Azab, H., Hayden, B., and Soltani, A. (2018). On the flexibility of basic risk attitudes in monkeys. Journal of Neuroscience. https://doi.org/10.1523/JNEUROSCI.2260-17.2018

Farashahi, S., Donahue, C. H., Hayden, B. Y., Lee, D., and Soltani, A. (2019). Flexible combination of reward information across primates. Nat. Hum. Behav., 3, 1215–1224.

Farovik, A., Place, R. J., McKenzie, S., Porter, B., Munro, C. E., and Eichenbaum., H. (2015). Orbitofrontal cortex encodes memories within value-based schemas and represents contexts that guide memory retrieval. The Journal of Neuroscience, 35, 8333–8344.

FitzGerald, T. H. B., Friston, K. J., and Dolan, R. J. (2012). Action-specific value signals in reward-related regions of the human brain. The Journal of Neuroscience, 32, 16417–16423.

FitzGerald, T. H. B., Seymour, B., Dolan, R. J. (2009). The role of human orbitofrontal cortex in value comparison for incommensurable objects. The Journal of Neuroscience, 29, 8388–8395.

Gallego, J. A., Perich, M. G., Miller, L. E., and Solla, S. A. (2017). Neural manifolds for the control of movement. Neuron, 94.

Gardner, M. P. H. and Schoenbaum, G. (2020). The orbitofrontal cartographer. Psyarxiv.

Glimcher, P. W., Dorris, M. C., and Bayer, H. M. (2005). Physiological utility theory and the neuroeconomics of choice. Games and Economic Behavior, 52, 213–256.

Gross, J., Woelbert, E., Zimmermann, J., Okamoto-Barth, S., Riedl, A. and Goebel, R. (2014). Value signals in the prefrontal cortex predict individual preferences across reward categories. Journal of Neuroscience, 34, 7580–7586.

Hayden, B. Y. (2017). Economic choice: The foraging perspective. Current Opinion in Behavioral Sciences, 24, 1–6.

Hayden, B., Heilbronner, S., and Platt, M. (2010). Ambiguity aversion in rhesus macaques. Frontiers in Neuroscience, 4.

Hayden, B. Y. and Niv, Y. (2020). The case against economic values in the brain. PsyArXiv.

Heilbronner, S. R. (2017). Modeling risky decision-making in nonhuman animals: shared core features. Current Opinion in Behavioral Sciences. https://doi.org/10.1016/j.cobeha.2017.03.001

Heilbronner, S. R. and Hayden, B. Y. (2013). Contextual factors explain risk-seeking preferences in rhesus monkeys. Frontiers in Neuroscience. https://doi.org/10.3389/fnins.2013.00007

Kable, J. K. and Glimcher, P. W. (2007). The neural correlates of subjective value during intertemporal choice. Nature Neuroscience, 10(12).

Kable, J. W. and Glimcher, P. W. (2009). The neurobiology of decision: Consensus and controversy. Neuron, 63, 733–745.

Kahnt, T., Heinzle, J., Park, S. Q., and Haynes, J-D. (2010). The neural code of reward anticipation in human orbitofrontal cortex. Proceedings of the National Academy of Sciences, 107, 6010–6015.

Kaufman, M. T., Churchland, M. M., Ryu, S. I., and Shenoy, K. V. (2015). Vacillation, indecision and hesitation in moment-by-moment decoding of monkey motor cortex. eLife. doi:10.7554/eLife.04677.001

Kennerley, S. W. & Wallis, J. D. (2009). Reward-dependent modulation of working memory in 689 lateral prefrontal cortex. Journal of Neuroscience, 29(10).

Kim, S., Bobeica, I., Gamo, N. J., Arnsten, A. F. T., and Lee, D. (2012). Effects of α-2A adrenergic receptor agonist on time and risk preference in primates. Psychopharmacology, 219, 363–375.

Klein, J. T., Deaner, R. O., and Platt, M. L. (2008). Neural correlates of social target value in macaque parietal cortex. Current Biology, 18, 419–424.

Knutson, B., Delgado, M. R., and Phillips, P. E. M. (2009). Chapter 25 - Representation of subjective value in the striatum. Neuroeconomics, Academic Press, 389–406.

Glimcher, P. W., Fehr, E., and Poldrack, R. A. (Ed.s). Krajbich, I., Armel, C., and Rangel A. (2010). Visual fixations and the computation and comparison of value in simple choice. Nature Neuroscience, 13, 1292–1298.

Lau, B. and Glimcher, P. W. (2008). Value representations in the primate striatum during matching behavior. Neuron, 58, 451–463.

Levy, D. J. and Glimcher, P. W. (2012). The root of all value: A neural common currency for choice. Current Opinion in Neurobiology. https://doi.org/10.1016/j.conb.2012.06.001

Lopes, L. L. (1987). Between hope and fear: The psychology of risk. Advances in sExperimental Social Psychology. https://doi.org/10.1016/S0065-2601(08)60416-5

Loewenstein, G. F., Weber, E. U., Hsee, C. K., and Welch, N. (2001). Risk as feelings. Psychological Bulletin, 127.

Lucantonio, F., Gardner, M. P. H., Mirenzi, A., Nerman, L. E., Takahashi, Y. K., and Schoenbaum, G. (2015). Neural estimates of imagined outcomes in basolateral amygdala depend on orbitofrontal cortex. Journal of Neuroscience, 35, 16521–16530.

Maisson, D. J., Cash-padgett, T. V, and Hayden, B. Y. (2020). A functional hierarchy for choice in medial prefrontal cortex. BioRxiv.

Marshall, J. A. R., Bogacz, R., Dornhaus, A., Planque, R., Kovacs, T., and Franks, N. R. (2009). On optimal decision-making in brains and social insect colonies. Journal of the Royal Society, Interface. 61065–1074

McCoy, A. N. and Platt, M. L. (2005). Risky-sensitive neurons in macaque posterior cingulate cortex. Nature Neuroscience, 8.

McGinty, V. B., Rangel, A., & Newsome, W. T. (2016). Orbitofrontal cortex value signals depend on fixation location during free viewing. Neuron, 90, 1299–1311.

Mehta, P. S., Tu, J.,C. LoConte, G. A., Pesce, M. C., and Hayden, B. Y. (2019). Ventromedial prefrontal cortex tracks multiple environmental variables during search. Journal of Neuroscience, 39, 5336–5350.

Montague, P. R. and Berns, G. S. (2002). Neural economics and the biological substrates of valuation. Neuron. https://doi.org/10.1016/S0896-6273(02)00974-1

Myers-Schulz, B. and Koenigs, M. (2012). Functional anatomy of ventromedial prefrontal cortex: implications for mood and anxiety disorders. Molecular Psychiatry, 17, 132–141.

Neubert, F-X., Mars, R. B., Sallet, J., and Rushworth, M. F. S. (2015). Connectivity reveals relationship of brain areas for reward-guided learning and decision making in human and monkey frontal cortex. PNAS, 112, E2695–E2704.

Noonan, M. P., Mars, R. B., and Rushworth, M. F. S. (2011). Distinct roles of three frontal cortical areas in reward-guided behavior. Journal of neuroscience, 31, 143999–14412.

O’Doherty, J., Kringelbach, M., Rolls, E. et al. (2001). Abstract reward and punishment representations in the human orbitofrontal cortex. Nature Neuroscience, 4, 95–102.

O’Donoghue, Ted, and Matthew Rabin. 2015. Present bias: Lessons learned and to be learned. American Economic Review, 105.

Oşan R., Zhu, L., Shoham, S., and Tsien, J. Z. (2007) Subspace projection approaches to classification and visualization of neural network-level encoding patterns. PLOS ONE, 2.

Padoa-Schioppa, C. (2011). Neurobiology of economic choice: A good-based model. Annual Review of Neuroscience, 3.

Padoa-Schioppa, C. and Assad, J. A. (2006). Neurons in the orbitofrontal cortex encode economic value. Nature. https://doi.org/10.1038/nature04676

Padoa-Schioppa, C. and Conen, K. E. (2017). Orbitofrontal cortex: A neural circuit for economic decisions. Neuron, 96, 736–754.

Padoa-Schioppa, C. and Schoenbaum, G. (2015). Dialogue on economic choice, learning theory, and neuronal representations. Current Opinion in Behavioral Sciences. https://doi.org/10.1016/j.cobeha.2015.06.004

Paxinos, G., Petrides, M., Huang, X., & Toga, A. W. (2008). The rhesus monkey brain in stereotaxic coordinates. Elsvier Science.

Pirrone, A., Azab, H., Hayden, B. Y., Stafford, T., & Marshall, J. A. R. (2018). Evidence for the speed–value trade-off: Human and monkey decision making is magnitude sensitive. Decision, 5(2), 129–142.

Platt, M. L. and Huettel, S. A. (2008). Risky business: The neuroeconomics of decision making under uncertainty. Nature Neuroscience, 11.

Rangel, A., Camerer, C., and Montague, P. (2008). A framework for studying the neurobiology of value-based decision making. Nature Review Neuroscience, 9, 545–556.

Rich, E. and Wallis, J. (2016). Decoding subjective decisions from orbitofrontal cortex. Nature Neuroscience, 19, 973–980.

Rolls, E. T. (2000). The orbitofrontal cortex and reward. Cerebral Cortex, 10, 284–294.

Rudebeck, P. H. and Murray, E. A. (2014). The orbitofrontal oracle: Cortical mechanisms for the prediction and evaluation of specific behavioral outcomes. Urology. https://doi.org/10.1016/j.neuron.2014.10.049

Rudebeck, P. H., Saunders, R. C., Lundgren, D. A., and Murray, E. A. (2017). Specialized representations of value in the orbital and ventrolateral prefrontal cortex: Desirability versus availability of outcomes. Neuron. https://doi.org/10.1016/j.neuron.2017.07.042

Rushworth, M. F. S., Noonan, M. A. P., Boorman, E. D., Walton, M. E., and Behrens, T. E. (2011). Frontal cortex and reward-guided learning and decision-making. Neuron, 70, 1054–1069.

Rustichini, A. and Padoa-Schioppa, C. (2015). A neuro-computational model of economic decisions. Journal of Neurophysiology, 114, 1382–1398.

Schoenbaum, G., Chiba, A. A., & Gallagher, M. (1998). Orbitofrontal cortex and basolateral amygdala encode expected outcomes during learning. Nature Neuroscience, 1, 155–159.

Schoenbaum, G., Setlow, B., Saddoris, M. R., and Gallagher, M. (2003). Encoding predicted outcome and acquired value in orbitofrontal cortex during cue sampling depends upon input from basolateral amygdala. Neuron, 39, 855–867.

Schoenbaum, G., Takahashi, Y., Liu, T. L., and Mcdannald, M. A. (2011). Does the orbitofrontal cortex signal value? Annals of the New York Academy of Sciences. https://doi.org/10.1111/j.1749-6632.2011.06210.x

Seeley, T.D., Camazine, S., and Sneyd, J. (1991). Collective decision-making in honey bees: how colonies choose among nectar sources. Behav Ecol Sociobiol 28, 277–290.

Seeley, T. D., Visscher, P. K., Schlegel, T., Hogan, P. M., Franks, N. R., and Marshall, J. A. (2012). Stop signals provide cross inhibition in collective decision-making by honeybee swarms. Science, 6, 108–111

Semedo, J. D., Zandvakili, A., Machens, C. K., Yu, B. M., and Kohn, A. (2019). Cortical areas interact through a communication subspace. Neuron, 102, 249–259.

Schuck, N. W., Cai, M. B., Wilson, R. C., and Niv, Y. (2016). Human orbitofrontal cortex represents a cognitive map of states space. Neuron, 91, 1402–1412.

Schuck, N. W. and Niv, Y. (2019). Sequential replay of nonspatial task states in the human hippocampus. Science, 80(364).

Sleezer, B. J., Castagno, M. D., and Hayden, B. Y. (2016). Rule encoding in orbitofrontal cortex and striatum guides selection. Journal of Neuroscience, 36, 11223–11237.

So. N. Y. and Stuphorn, V. (2010). Supplementary eye field encodes option and action value for saccades with variable reward. Journal of Neurophysiology, 104, 2634–2653.

So, N. Y. and Stuphorn, V. (2016). Supplementary eye field encodes confidence in decisions under risk. Cerebral Cortex, 26, 764–782.

Stokes, M., Muhle-Karbe, P. S., Myers, N. E. (2020). Theoretical distinction between functional states in working memory and their corresponding neural states. PsyArXiv.

Strait, C. E., Blanchard, T. C., and Hayden, B. Y. (2014). Reward value comparison via mutual inhibition in ventromedial prefrontal cortex. Neuron, 82.

Strait, C. E., Sleezer, B. J., Blanchard, T. C., Azab, H., Castagno, M. D., and Hayden, B. Y. (2016). Neuronal selectivity for spatial positions of offers and choices in five reward regions. Journal of Neurophysiology. https://doi.org/10.1152/jn.00325.2015

Strait, C. E., Sleezer, B.J., and Hayden, B. Y. (2015). Signatures of value comparison in ventral striatum neurons. PLOS Biology, https://doi.org/10.1371/journal.pbio.1002173

Staudinger, M. R., Erk, S., Abler, B. and Walter, H. (2009). Cognitive reappraisal modulates expected value and prediction error encoding in the ventral striatum. NeuroImage, 47, 713–721.

Takahashi, Y. K., Roesch, M. R., Wilson, R. C., Toreson, K., O’Donnell, P., Niv, Y., and Schoenbaum, G. (2011). Expectancy-related changes in firing of dopamine neurons depend on orbitofrontal cortex. Nature Neuroscience. https://doi.org/10.1038/nn.2957

Tremblay, L. and Schultz, W. (1999). Relative reward preference in primate orbitofrontal cortex. Nature, 398, 704–708.

Vlaev, I., Chater, N., Stewart, N., and Brown, G. D. A. (2011). Does the brain calculate value? Trends in Cognitive Sciences. https://doi.org/10.1016/j.tics.2011.09.008

Wallis, J. D. (2007). Orbitofrontal cortex and its contribution to decision-making. Annu. Rev. Neurosci, 30, 31–56

Wang, M. and Hayden, B. (2017). Reactivation of associative structure specific outcome responses during prospective evaluation in reward-based choices. Nature Communications, 8. https://doi.org/10.1038/ncomms15821

Wikenheiser, A. M. and Schoenbaum, G. (2016). Over the river, through the woods: cognitive maps in the hippocampus and orbitofrontal cortex. Nature Reviews Neuroscience, 17, 513–523.

Wilson, R. C., Takahashi, Y. K., Schoenbaum, G., and Niv, Y. (2014). Orbitofrontal cortex as a cognitive map of task space. Neuron, 81, 267–279.

Wunderlich, K., Dayan, P., and Dolan, R. (2012). Mapping value based planning and extensively trained choice in the human brain. Nature Neuroscience, 15, 786–791.

Xie, J. and Padoa-Schioppa, C. (2016). Neuronal remapping and circuit persistence in economic decisions. Nature Neuroscience, 19, 855–861.

Xie, Y., Nie, C., & Yang, T. (2018). Covert shift of attention modulates the value encoding in the orbitofrontal cortex. eLife, 7, e31507.

Yoo, S. B. M. and Hayden, B. Y. (2018). Economic choice as an untangling of options into actions. Neuron, 99(3), 434–447. https://doi.org/10.1016/j.neuron.2018.06.038

Yoo, S. B. M. and Hayden, B. Y. (2020). The transition from evaluation to selection involves neural subspace reorganization in core reward regions. Neuron, 105.

